# Murine models of renal ischaemia reperfusion injury: An opportunity for refinement using non-invasive monitoring methods

**DOI:** 10.1101/2019.12.17.879742

**Authors:** Rachel Harwood, Joshua Bridge, Lorenzo Ressel, Lauren Scarfe, Jack Sharkey, Gabriella Czanner, Philip Kalra, Aghogho Odudu, Simon Kenny, Bettina Wilm, Patricia Murray

**Affiliations:** Institute of Translational Medicine, University of Liverpool; Alder Hey Children’s Hospital, Liverpool; Department of Biostatistics, University of Liverpool; Department of Veterinary Pathology and Public Health, University of Liverpool; Division of Cardiovascular Sciences, University of Manchester; Salford Royal NHS Foundation Trust, Salford

**Keywords:** Acute Kidney Injury, Chronic Kidney Disease, Ischaemia Reperfusion Injury

## Abstract

**Background:** Renal Ischaemia Reperfusion Injury (R-IRI) can cause Acute Kidney Injury (AKI) and Chronic Kidney Disease (CKD), resulting in significant morbidity and mortality. To understand the underlying mechanisms, reproducible small-animal models of AKI and CKD are needed. We describe how innovative technologies for measuring kidney function non-invasively in small rodents allow successful refinement of the R-IRI models, and offer the unique opportunity to monitor longitudinally in individual animals the transition from AKI to CKD.

**Methods:** Male BALB/c mice underwent bilateral renal pedicle clamping (AKI) or unilateral renal pedicle clamping with delayed contralateral nephrectomy (CKD) under isoflurane anaesthetic. Transdermal GFR monitoring and multi-spectral optoacoustic tomography in combination with statistical analysis were used to identify and standardise variables within these models.

**Results:** Pre-clamping anaesthetic time was one of the most important predictors of AKI severity after R-IRI. Standardising pre-clamping time resulted in a more predictably severe AKI model. In the CKD model, initial improvement in renal function was followed by significant progressive reduction in function between weeks 2 and 4. Performing contralateral nephrectomy on day 14 enabled the development of CKD in a survivable way.

**Conclusions:** Non-invasive monitoring of global and individual renal function after R-IRI is feasible, reproducible and correlates well with classical markers of injury. This facilitates refinement of kidney injury models and enables the degree of injury seen in pre-clinical models to be translated to those seen in the clinical setting. Thus, future therapies can be tested in a clinically relevant, non-invasive manner.

**What is already known:** The severity of Renal Ischaemia Reperfusion injury (R-IRI) varies between animal strain, gender and age. Experimental variables including temperature and clamping time are usually tightly controlled but significant variability still exists. Classically, small rodent experiments depend on endpoint evaluation of serum and histological features of disease. However, new technologies including transdermal glomerular filtration rate (GFR) monitoring and Multispectral Optoacoustic Tomography (MSOT) may enable renal function to be accurately monitored longitudinally, enabling better refinement of these models.

**What this study adds:** This study shows that transdermal GFR measurements have reliably enabled refinement of the R-IRI model by standardisation of the duration of isoflurane prior to commencing surgery. Individual kidney function can be assessed in-vivo after unilateral R-IRI using MSOT imaging. The excretion t_max_ of IRDye-800 reliably represents the relative function of the injured kidney, permitting longitudinal in-vivo assessment of differential kidney function.

**What impact this may have on practice:** This study demonstrates the utility of two minimally-invasive in-vivo methods of monitoring kidney function which have advantages over classical methods and potentially enable fewer animals to be used in future studies. The study demonstrates refinement of bilateral and unilateral R-IRI models which will also enable a reduction in the number of animals needed for experimentation.

## Introduction

Acute kidney injury (AKI) and chronic kidney disease (CKD) are leading causes of morbidity and mortality^1^. Pre-renal causes of AKI can be attributed to hypovolaemia^2^, cardiac bypass surgery^3^ and clamping of the renal pedicle during partial nephrectomy^4^. These lead to renal ischaemia-reperfusion injury (R-IRI) which progresses from AKI to functional recovery in most^2^, although severe R-IRI AKI is a well-recognised cause of CKD. AKI in children significantly increases their risk of developing CKD^5,6^ and an episode of AKI in an adult with CKD accelerates their progression towards end-stage renal disease (ESRD)^7^.

Despite the need for therapeutic adjuncts to reduce the severity of AKI and progression to CKD, therapies are still lacking. A range of potential treatments have been investigated, but none are recommended due a lack of efficacy in clinical trials^8^. In view of this on-going need, robust pre-clinical evaluation of novel therapies for AKI and CKD is required.

R-IRI is a pre-clinical model of AKI^9^ caused by unilateral or bilateral clamping of the renal pedicles, and in the former, with or without contralateral nephrectomy. Creation of R-IRI varies widely between groups who use differing anaesthetic agents, clamping times, temperature control and approaches to the kidney^10-12^, all of which could cause alteration in the severity of R-IRI induced injury. Additionally, the genetic background and strain of animal can affect the progression of AKI and CKD^13^. Because R-IRI is a notoriously variable model of kidney injury^12^, its refinement enables fewer animals to be used when assessing therapeutic effects.

Bilateral R-IRI creates an injury that affects the global glomerular filtration rate (GFR), akin to the clinical situations described above. A global reduction in GFR is classically measured using blood markers of kidney function which do not have a linear relationship with GFR^14,15^. In small rodents these are typically performed at the experimental endpoint due to the volumes of blood required^16^. By contrast, non-invasive methods of measuring renal clearance enable monitoring of kidney function in the same animal^17,18^. Transdermal measurement of FITC-Sinistrin clearance time allows the calculation of a global GFR^18,19^, making it an ideal tool for longitudinally monitoring renal function after bilateral R-IRI and reducing the numbers of animals required for a study.

The Kidney Disease Improving Global Outcomes (KDIGO) group released guidance in 2012^20^ to amalgamate previous AKI stratification tools, primarily the Acute Dialysis Quality Initiative (ADQI)^21^ who defined the ‘RIFLE’ classification and the Acute Kidney Injury Network (AKIN)^22^ guidance. The KDIGO guidance uses increase in serum Creatinine and urine output to define AKI. Pre-clinically, serial serum and urinary measurements are not possible in small rodents. The Acute Dialysis Quality Initiative (ADQI) group stratified AKI primarily using reduction in GFR. Levels of injury are classed as ‘Risk’ (>25% GFR decrease), ‘Injury’ (>50% GFR decrease), ‘Failure’ (>75% GFR decrease), ‘Loss’ (complete loss of function > 4 weeks) and ‘ESRD’^21^. In AKI the patients most in need of additional therapies are those in the ‘Failure’ group^23^ who have a 46.5% mortality rate. Developing a clinically relevant pre-clinical model of AKI enables therapies to be trialled appropriately^24^.

Unilateral R-IRI with delayed nephrectomy allows a more severe AKI to be induced because the healthy kidney provides function whilst the severely injured kidney recovers adequate function for survival. It is a model that enables both to assess the progression to CKD as well as the most severe manifestation of AKI. When a unilateral R-IRI is performed without nephrectomy the overall GFR does not reflect the function of the injured kidney, which will be substantially less than the uninjured kidney. This makes it difficult to establish the optimal time point for removing the healthy kidney to allow transdermal GFR monitoring of the injured kidney. From an NC3Rs perspective, refinement of this model is important because if the healthy kidney is removed before the injured kidney has regained sufficient function, the animal will experience suffering.

We have previously shown that measurements of IRDye 800 clearance^25^ using multispectral optoacoustic tomography (MSOT) correlate well with histological evidence of kidney damage in an Adriamycin model of injury^16^. Moreover, a recent study has shown that IRDye 800 can be measured repeatedly and reproducibly in the kidneys of healthy mice using MSOT, indicating that longitudinal studies could be performed with high confidence^26^. In unilateral R-IRI, non-invasive monitoring of both the injured and healthy kidney using MSOT could enable the function of each kidney to be simultaneously and individually tracked.

In this study we used a validated transcutaneous device to measure GFR longitudinally in mice, in order to establish the effect of clamping time on GFR in the bilateral R-IRI model^13,16^. This allowed us to determine whether GFR returned to baseline levels after a 2 week recovery period. It also enabled us to explore whether the degree of GFR impairment that occurs immediately following R-IRI, can predict the degree of histological damage that occurs subsequently. By exploring the contribution of anaesthetic time to the severity of injury, we refined the model to reduce variability. Finally, we developed a clinically relevant model of AKI, representing severe AKI leading to CKD, but remaining NC3Rs compliant. To this end, we explored the potential of an emerging technology, MSOT, which enables non-invasive visualisation of the kidneys, and monitoring of their function longitudinally.

## Materials and Methods

### Animals

All experimentation was performed under a Project Licence (PPL 7008741) granted by the Home Office under the Animals (Scientific Procedures) Act 1986. Male BALB/c mice (Charles River, Margate, UK) aged 8-10 weeks were used throughout. Mice were housed in individually ventilated cages in a 12 hour light/dark cycle. They had *ad libitum* access to food and water and were acclimatised for 1 week before entering the study. Experiments are reported in line with the ARRIVE guidelines and further information can be found in Supplementary Information 1.e.

### AKI Model

Briefly, after induction of anaesthesia with inhaled oxygenated isoflurane, the animal was transferred to a heat pad with a rectal temperature probe and a feedback-regulated system, body temperature set to 37°C. Surgery was commenced after the temperature reached 36.5°C. A dorsal approach was taken^27^ and an atraumatic vascular clamp (Schwartz, Interfocus, Linton) was placed on the vessels. Clamp times of 25, 27.5 and 30 minutes were used with 6 animals per group. For further details, see Supplementary Information 1.a and 1.b. Data were collected on the duration of anaesthesia before the first clamp was applied and the duration between the removal of the second clamp and the mouse entering the post-operative warmed chamber (post-clamp time). Three mice underwent sham operations. Animals underwent FITC-Sinistrin clearance measurement pre-operatively and on days 1, 3, 7 and 14 after surgery. On day 14, animals were culled and blood and kidneys collected.

### Refined AKI Model

A refined AKI model was performed in 6 animals. The main adjustment was standardisation of pre-clamping anaesthetic time to 30 minutes between the start of anaesthetic and commencement of surgery. For further details see Supplementary Information 1.c and Supplementary Figure 1. GFR was measured pre-operatively and day 1 and 3 post- operatively. Blood and kidneys were collected after animals were culled on day 3.

### CKD Model

To cause CKD secondary to R-IRI, prolonged unilateral renal pedicle clamping was performed in 3 mice using the ‘Refined AKI model’ with clamping of the left renal pedicle for 40 minutes. MSOT was undertaken on days 1, 4, 7, 10, 14, 21 and 28 after injury, and nephrectomy of the uninjured kidney was performed on day 29 (Supplementary Figure 2A). GFR was measured on day 30, animals were culled, and blood and kidneys retrieved.

### Refined CKD Model

Following the first CKD experiment, the model was refined by performing right-sided renal pedicle clamping in 6 animals, enabling better visualisation of the kidney with MSOT. The contralateral nephrectomy was performed on day 14 and the endpoint of the experiment was 6 weeks after R-IRI (Supplementary Figure 2B).

### Transdermal FITC-Sinistrin Clearance Measurement

FITC-Sinistrin clearance was measured in all mice, as previously described^19^. The half-life of

FITC-Sinistrin was converted to GFR using the formula^18^

GFR [μl/min/100g bw] = 14616.8 [μl/100g bw]/t_1/2_(FITC-Sinistrin) [min] to enable the GFR of mice with no excretion to be included in the analysis.

Absolute GFR values and the proportional change in GFR compared to baseline measurements were analysed. The proportional change was correlated with the ‘RIFLE’ classification of kidney injury, depicting the levels of Risk, Injury and Failure as described by the ADQI working group^21^. The values for each level were calculated from the mean of the baseline measurements, which followed a normal distribution (Supplementary Figure 3).

### Multi-Spectral Optoacoustic Tomography (MSOT)

The inVision 256-TF MSOT imaging system (iThera Medical, Munich, Germany) was employed in the CKD model, using IRDye 800 to measure renal kinetics in the injured and healthy contralateral kidneys (details of acquisition and analysis within Supplementary Information 1.d).

### Serum analysis

Animals were culled in a rising concentration of CO_2_. Once death was confirmed, cardiac puncture was performed to collect blood. This was clotted for 2 hours and centrifuged at 2000g for 20 minutes. Serum was collected and re-centrifuged at 75g for 4 minutes to remove all blood. Samples were stored at -20°C until analysis and repeat freeze-thawing was avoided. Creatinine (Detect X Serum Creatinine Detection Kit, Arbor Assays, Ann Arbor, MI, USA) was assessed using the modified Jaffe Reaction. Urea (QuantiChrom Urea Assay Kit, BioAssay Systems, Hayward, CA, USA) was assessed using a colorimetric reaction and Cystatin C (Quantikine ELISA, R&D Systems, UK) was analysed using an ELISA method.

### Histopathology

Kidneys were collected and fixed in 4% paraformaldehyde for 1 day. They were processed in paraffin and sections of 4 µm were stained with and Picrosirius Red (PSR) to assess for cortical deposition of collagen I and III. Sections were analysed by a veterinary histopathologist blinded to the intervention. Cortical collagen deposition was graded 0-4 in a single section of each kidney, based on the percentage of the cortex judged to show increased collagen deposition by the histopathologist (Supplementary Figure 6).

### Statistical Analysis

One-way ANOVA was used to compare GFR at different time points after injury, serum and histological results. A mixed effects model was used to describe longitudinal weight data of all surviving animals and to analyse t_max_ results. A post-analysis Bonferroni correction for comparisons between all groups was applied. Linear regression analysis compared GFR with other variables including anaesthetic time, heat pad and surgical order and serum and histological markers of kidney injury. Significance was taken as p<0.05.

## Results

### In bilateral R-IRI, GFR is inversely correlated with subsequent collagen deposition

Bilateral R-IRI was performed using 3 different clamp times – 25, 27.5 and 30 min – to establish which was best able to induce a severe, survivable injury. Compared with the sham group, the GFR of all injured animals was significantly reduced on day 1, irrespective of clamp time, but had returned to near baseline levels by day 14. There was no significant difference between R-IRI groups, but the trend was for the GFR to be lower in animals with longer clamp times (Figure 1A). The proportional GFR change showed that the majority of animals entered the RIFLE ‘failure’ category on day 1 after injury in the 27.5 and 30 min groups, but had mostly recovered by day 14 (Figure 1B).

**Figure 1.**
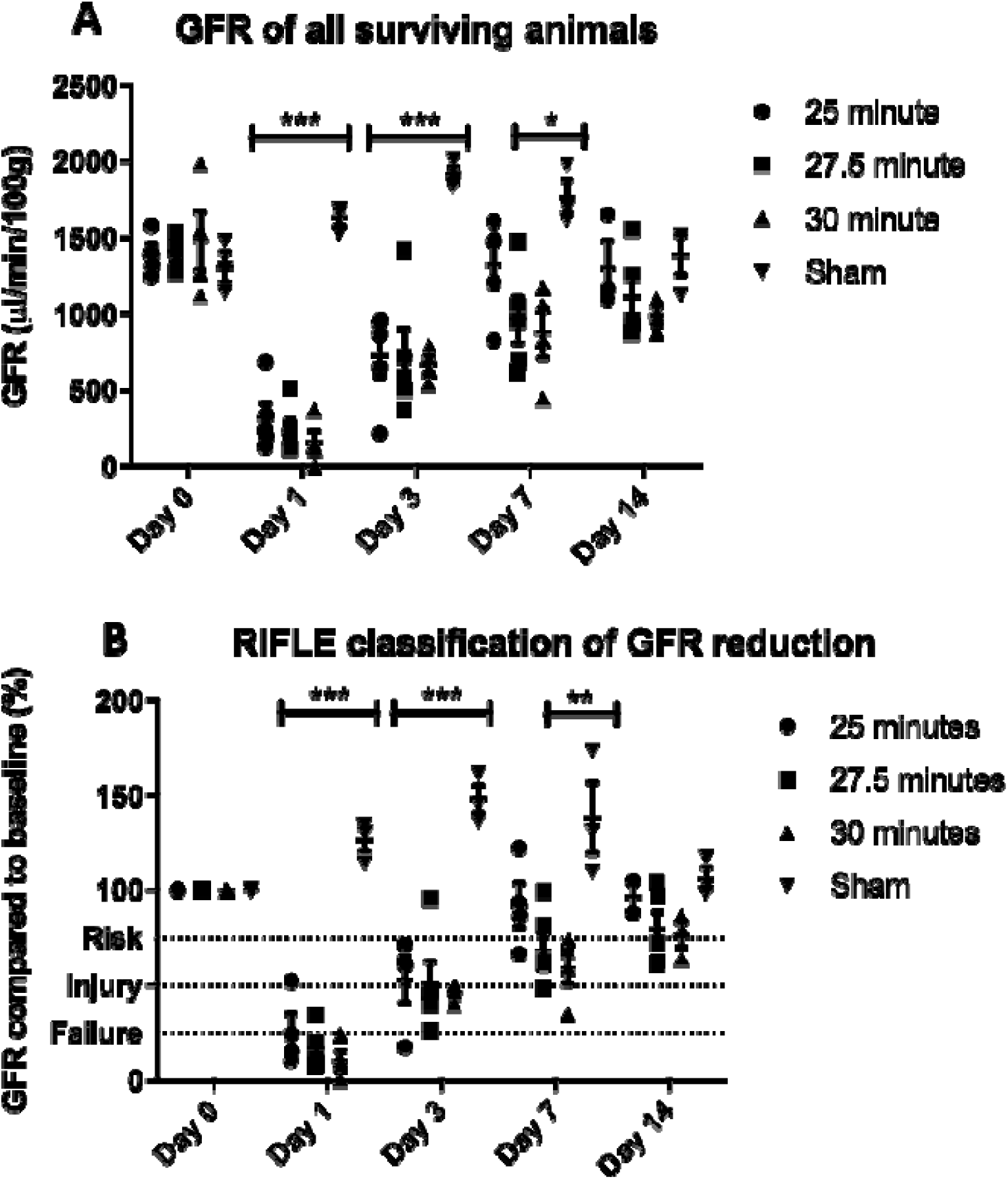
GFR results of animals undergoing bilateral R-IRI for 25, 27.5 and 30 minutes and sham procedures. A: Absolute change in GFR after R-IRI in animals who survived to the endpoint of the experiment. B: Proportional change in GFR after bilateral R-IRI. The ADQI ‘RIFLE’ criteria are shown within the figure. n=6:per clamped group, n=3:sham. All figures show mean ± SEM. * p < 0.05 compared to sham group, ** p < 0.01 compared to sham group, *** p < 0.001 compared to sham group.

There was weight loss in all groups with the nadir occurring 4 days after R-IRI (Supplementary Figure 4A). The mortality rate was higher in the 30 min group (33% compared with 17% in the 27.5 and 25 min groups; Supplementary Figure 4B). At the study end-point (day 14), no biologically relevant differences in serum creatinine, urea and cystatin-C were observed compared to sham controls (Figure 2). Because of its lower mortality rate than the 30 min group combined with a strong injury response at day 1 similar to the 30 min group, a clamp-time of 27.5 min was used in subsequent bilateral R-IRI experiments.

**Figure 2.**
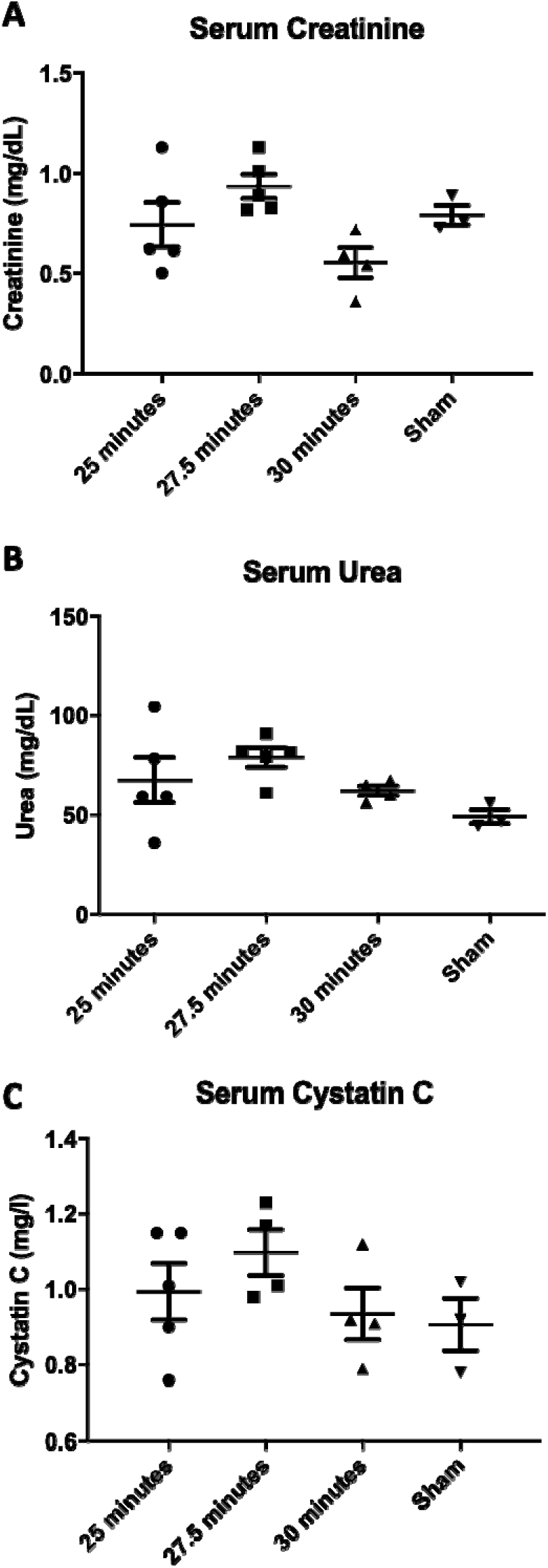
Serum analysis on day 14 after R-IRI. A: Serum creatinine, assayed using the modified Jaffe method. B: Serum Urea. C: Serum Cystatin C. There was no significant difference between the treatment groups for any of the parameters tested.

Our analysis also demonstrated that longer clamping was associated with increased collagen deposition on day 14 (Figure 3A), which showed a strong correlation with the day 1 GFR (Figure 3B). This indicates that following R-IRI, the risk of developing renal fibrosis is related to the degree of initial GFR impairment, reflecting the severity of the initial injury. A surprising finding, however, was that the GFR was not substantially affected by longer clamp times, raising the question of whether other extrinsic factors, such as anaesthetic time, could be affecting the severity of injury.

**Figure 3.**
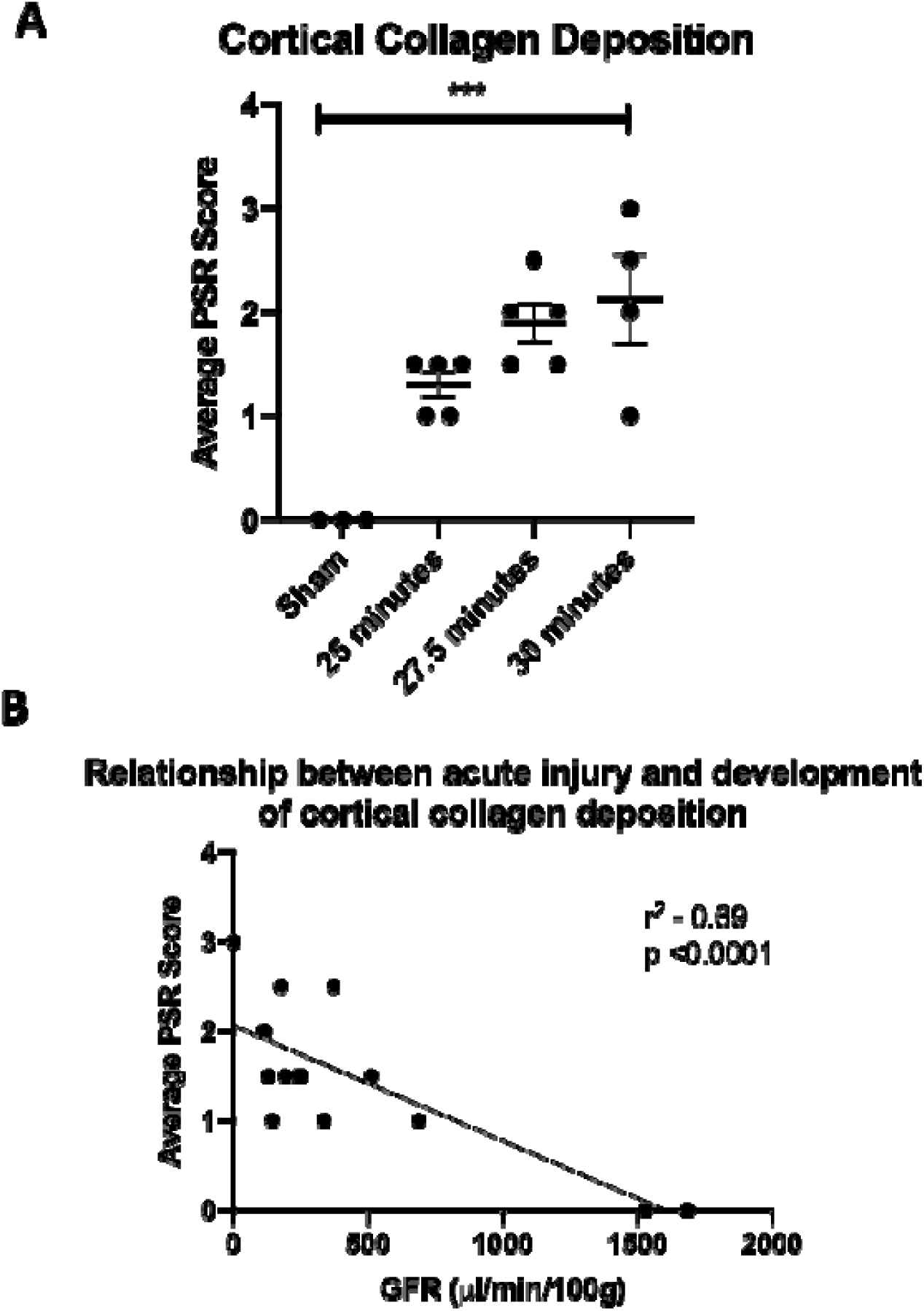
Cortical collagen deposition after R-IRI on day 14. A: Effect of clamping time on development of collagen deposition (Mean, SEM). Sham-operated animals had no collagen deposition. *** p < 0.001 compared to sham group when compared by one-way ANOVA. B: Association between the day 1 GFR and the average PSR score, showing that a more severe AKI leads to increased collagen deposition.

### Pre-clamping anaesthetic time correlates with the degree of GFR impairment

The model of bilateral R-IRI is notoriously variable^11^, even when clamp times and temperature are standardised. To determine the impact of external experimental variables on GFR, univariate linear regression analysis was performed. This revealed that, independent of clamping time, the total anaesthetic time had an impact on the day 1 GFR. When considering each component of the anaesthetic time, the GFR was particularly affected by the time between anaesthetic commencement and placement of first clamp (Figure 4A). There was no significant relationship between clamping time or post-clamping time and GFR (Figure 4B), nor surgical order, which heat pad was used, or the animal’s weight (Figure 4C). The total anaesthetic times were 78 minutes (Std Dev 13), 87 minutes (Std Dev 8) and 85 minutes (Std Dev 9) for the clamp times of 25, 27.5 and 30 minutes respectively. There is no significant difference between the total anaesthetic times for each group, showing that a longer clamping time did not equate to a longer anaesthetic time (p 0.40).

**Figure 4.**
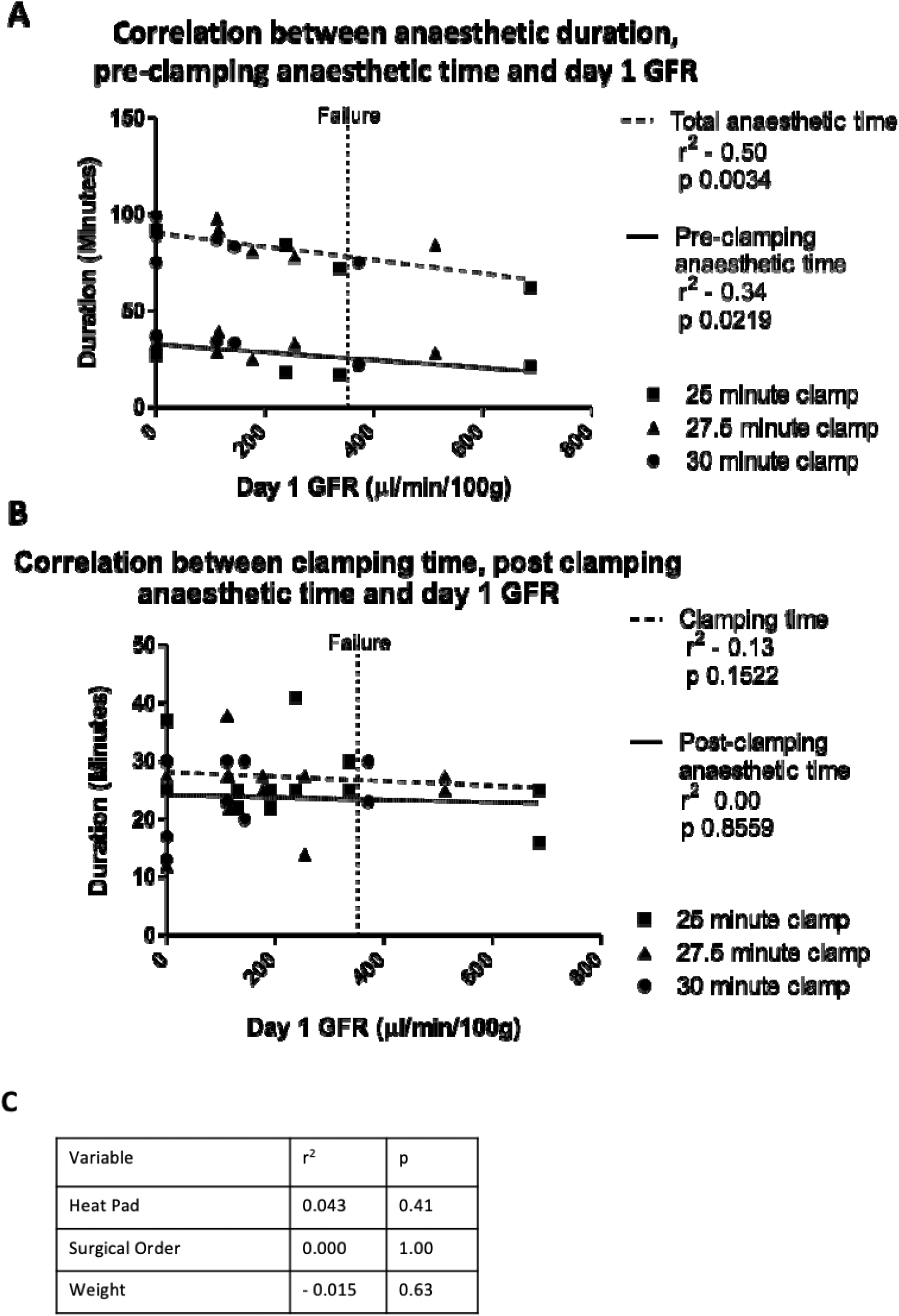
Impact of experimental variables on day 1 GFR measurement. A: Significant correlation of GFR with the total anaesthetic time and pre-clamping time. B: No correlation between GFR and clamping time or post-clamping time. C: No correlation between additional variables and GFR.

Based on these observations, we refined the bilateral R-IRI model (27.5 min clamp time) by standardising the pre-surgical anaesthetic time to 30 minutes from the start of anaesthetic to start of surgery, whilst continuing to monitor total anaesthetic duration. We compared this standardised anaesthetic (SA) group to the previously described non-standardised anaesthetic (NSA) group and found that the SA group had a reduction in day 1 GFR (Table 1), suggesting that a more severe but more reproducible injury level was achieved. Standardising pre-surgical time meant that surgery was more regimented, resulting in a significantly shorter overall anaesthetic time, but a longer, less variable pre-clamping time. This highlights that it is pre-clamping anaesthetic time, rather than total anaesthetic duration, that has the greatest impact on the GFR.

**Table 1.** Comparison of non-standardised and standardised pre-surgical anaesthetic time. The GFR is reduced to within the ‘Failure’ category (<353µl/min/100g) for all animals who had a standardised pre-surgery anaesthetic time (SA) compared to those who has a non-standardised anaesthetic time (NSA). The pre-clamping anaesthetic time is much more tightly controlled with a significantly smaller variance (p 0.02). Values are Mean (St Dev).

|  | Non-standardised<br>pre-surgical<br>anaesthetic time | Standardised<br>pre-surgical<br>anaesthetic time |
| --- | --- | --- |
| Day 1 GFR<br>(μl/min/100g) | 195 (176.6) | 113 (121.3) |
| Total anaesthetic<br>time (min) | 86.60 (8.2) | 71.67 (9.5) |
| Pre-clamping<br>anaesthetic time<br>(min) | 30.8 (5.4) | 34.0 (1.5) |

### Assessing individual renal function in a mouse model of R-IRI-induced CKD using MSOT

A limitation of the bilateral R-IRI model is that it does not model the most severe AKI which consistently lead to CKD. Whilst patients with severe AKI can be kept alive with dialysis until renal recovery, mice cannot and very severe AKI is fatal. This can be overcome by prolonged clamping of one kidney to induce a very severe injury, and then removing the healthy kidney after the injured kidney has recovered. If the injured kidney has not recovered sufficiently before contralateral nephrectomy there is potential for animal suffering. To address this, we used MSOT to establish the time required for sufficient recovery of function. We first performed a pilot experiment with 40 min unilateral clamping, and monitored single kidney function in both the healthy and injured kidney with MSOT over 4 weeks by longitudinally assessing the pharmacokinetics (PK) of IRDye in the renal cortex (Supplementary Figure 2).

We established the most appropriate model for monitoring IRDye PK by testing the fit of bi-exponential and tri-exponential models to MSOT curves. Tri-exponential modelling fitted better than other models, both visually (Supplementary Figure 5) and using the Akaike Information Criterion^28^. The *t* _*max*_(time between administration of IRDye and maximum peak of the curve) distinguished between injured and uninjured kidneys consistently and was therefore further tested using linear mixed effects models. Using the uninjured kidney as a reference group there were significant differences between injured and uninjured kidney (p=0.044) on day 1 (Figure 5A). The injured kidney’s function improved to levels equivalent to the uninjured kidney in week 2 (p=0.75) but then declined significantly over the subsequent 2 weeks (Figure 5B).

**Figure 5.**
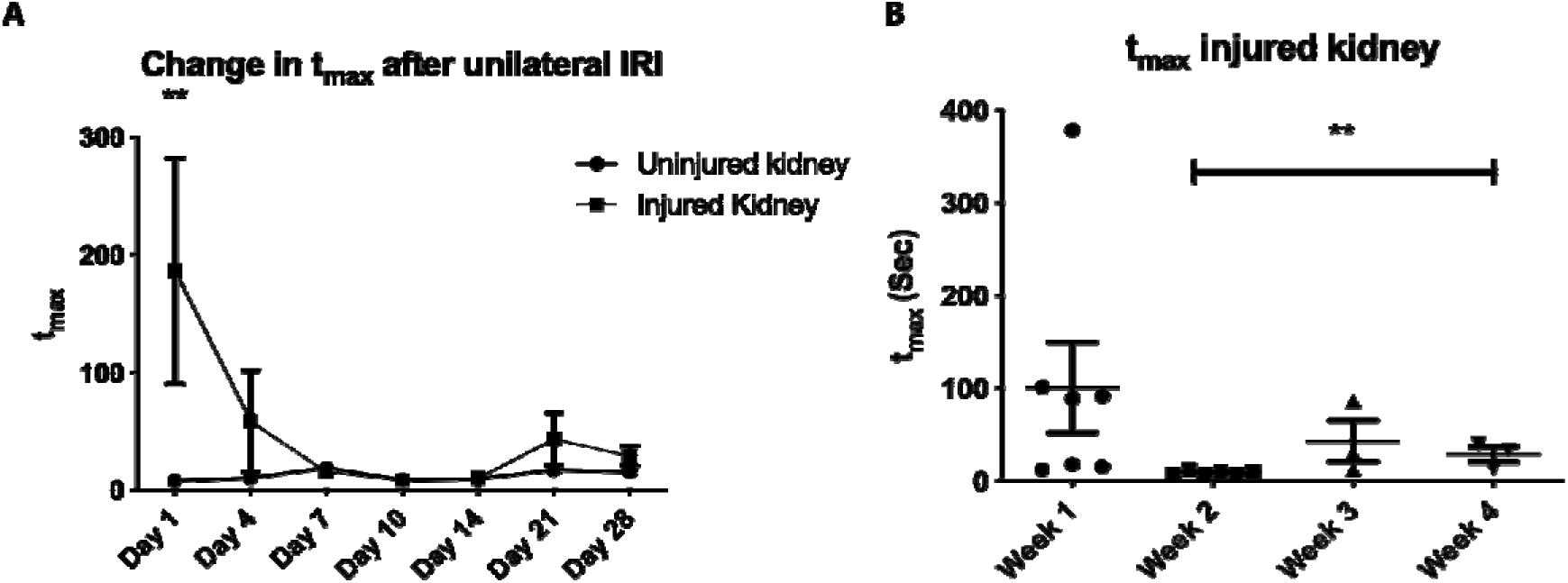
MSOT analysis of renal function in the CKD model. A: Change in t_max_ of IRDye-800 for the injured and uninjured kidney after IRI. On day 1 after injury there is a significant difference in function between the kidneys. B: The change in t_max_ of the injured kidney in the weeks after injury. There is a significant prolongation in t_max_ between week 2 and week 4 in the injured kidney. n=3. ** p < 0.01.

GFR measurements performed at the endpoint, one day after nephrectomy on day 29, revealed significant reduction in GFR to an average of 20% of baseline function (Figure 6A), showing progression from AKI to CKD. Serum markers of function reflected significant injury above published normal ranges (Supplementary Table 2)^29,30^. All injured kidneys weighed less than the uninjured kidneys, and there was collagen deposition within all injured kidneys (Figure 6B,C). The impaired GFR at the study end-point, accompanied with increased levels of serum biomarkers and histological damage, indicated that 40 min clamping with subsequent nephrectomy produces a severe renal injury resulting in CKD.

**Figure 6.**
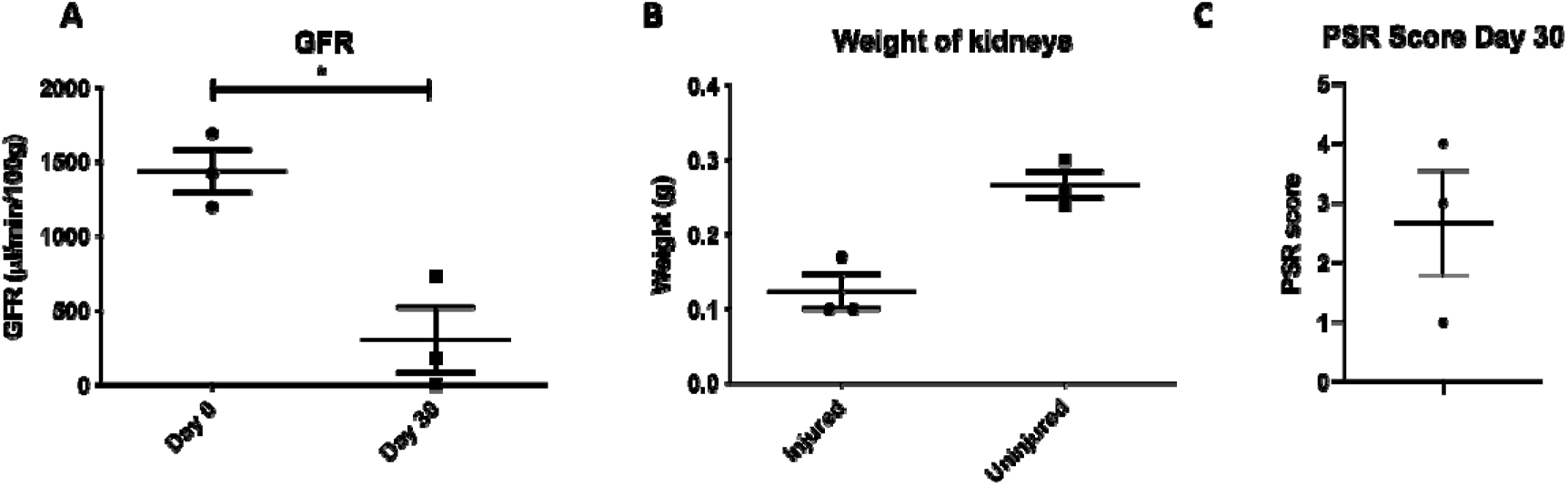
Renal injury in the clamped kidney after nephrectomy. A: GFR prior to IRI injury vs after R-IRI and contralateral nephrectomy shows significant reduction in GFR. B: The injured kidney weighs less than the uninjured kidney, showing loss of renal mass after prolonged R- IRI. C: All injured kidneys displayed histological evidence of collagen deposition when stained with Picrosirius Red (PSR). No collagen deposition is seen in uninjured kidneys.

### Refinement of a unilateral model of R-IRI-induced CKD results in a survivable, reproducible and persistent renal injury

In the pilot study of unilateral R-IRI and subsequent contralateral nephrectomy, MSOT showed a nadir of injury between days 10 and 14 (Figure 5A). Performing contralateral nephrectomy when the injured kidney is functioning optimally reduces animal suffering and improves survival. It also enables transdermal FITC-Sinistrin clearance to be measured, which is quicker to perform than MSOT and has the advantage that it can be undertaken in awake animals^16,31^. Therefore, the CKD model was refined by performing contralateral nephrectomy on day 14 after R-IRI (Supplementary Figure 2B).

In the refined study, the t_max_ differentiated between the injured and uninjured kidney on day 1 after R-IRI and significant reduction in t_max_ was seen between day 1 and day 9 after R- IRI (Figure 7A) and was maintained between day 9 and 13. One animal developed diarrhoea on day 12 and was excluded from further experimentation. Following contralateral nephrectomy on day 14, all animals had significant, persistent reduction in GFR (Figure 7B), without any significant changes in weight (Figure 7C). All animals had elevated serum markers of kidney injury (Supplementary Table 1) and consistent collagen deposition (Figure 7D). This demonstrates that MSOT is suitable to monitor individual kidney function after R- IRI using the t_max_ of IRDye clearance, and that doing so has enabled a tightly controlled model of AKI to CKD progression.

**Figure 7.**
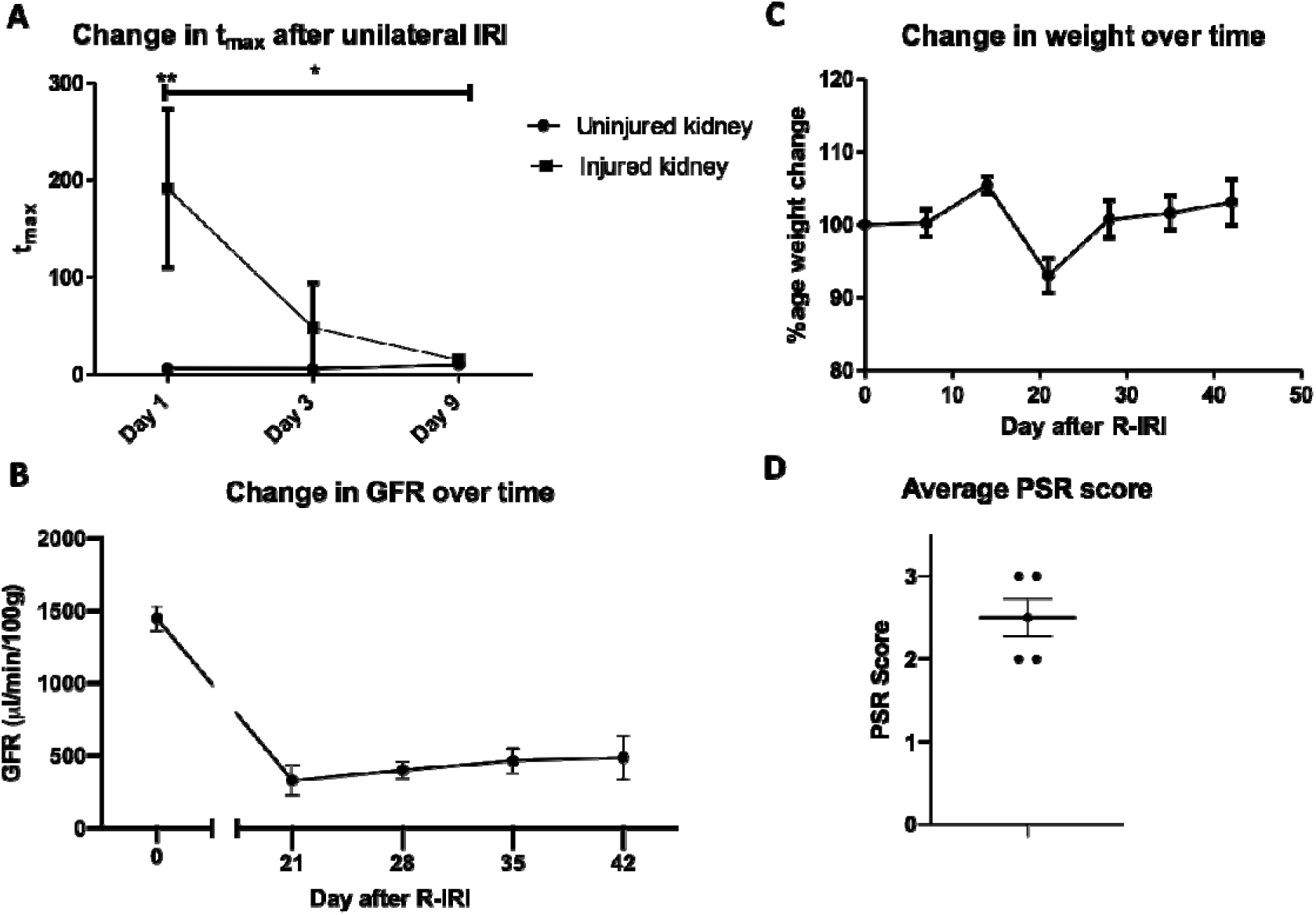
Refined CKD model. A: The t_max_ differentiates between the injured and uninjured kidney on day 1. There is significant reduction in t_max_ after injury, showing recovery of the kidney. * p < 0.05, ** p < 0.01. B: There is a persistent and significant reduction in GFR after contralateral nephrectomy. C: There is no significant change in weight. D: All animals had consistent levels of collagen deposition after staining with PSR. n=6 in A; n=5 in B-D.

Overall, we show that performing a contralateral nephrectomy on day 14 after R-IRI caused consistent, survivable CKD which can be monitored longitudinally with transdermal FITC-Sinistrin clearance.

## Discussion

Here we report, for the first time, how non-invasive monitoring of renal function can be used to identify factors which add variability to the R-IRI model, leading to its refinement. R- IRI represents a variety of clinical situations^9^ and its applicability to clinical practice is becoming increasingly evident with the recognition that one episode of ischaemic AKI can lead to nephron loss, tubule-interstitial atrophy and fibrosis.

The majority of clinical causes of R-IRI AKI affect both kidneys and therefore a bilateral R-IRI model was selected for AKI studies. Clinical literature describes that a severe episode of AKI is associated with an increased risk of developing CKD^32^ and the finding of collagen deposition 14 days after R-IRI AKI supports this. Concluding AKI studies on day 3 prevented the most severe weight loss in animals whilst still enabling the study of acute derangement of renal function. However, extending the study period enables the transition of AKI to CKD to be further examined. When considering CKD it was apparent that a survivable but severe, consistent model of kidney injury was required. Prolonged unilateral clamping with delayed nephrectomy caused less suffering to the animal but a consistent, reproducible reduction in GFR with more collagen deposition and less variability. When selecting whether unilateral or bilateral R-IRI be undertaken, the suffering of animals needs to be taken into account. The loss of animals can have an effect on experimental results and this should be considered when planning the experiments.

Description of the GFR as both an absolute measure and as a proportional measure of the baseline GFR enables the RIFLE classification^21^ to be applied to assess loss of renal function. This approach to describing the GFR is novel and enables results to be reported using clinically relevant descriptors. Therefore, applying RIFLE enables appropriate power calculations to be performed for therapeutic studies.

### Standardisation of pre-clamping anaesthetic time achieves a more reproducible R-IRI model

Refinement of the R-IRI model of AKI was possible because the duration of anaesthetic prior to clamping has a significant impact on day 1 GFR. Older studies have suggested that isoflurane is renoprotective^33^ but recent work disputes these findings^34^. In our experiments, we found that isoflurane did not prevent severe injury after R-IRI. We postulate that the duration of isoflurane anaesthetic is important due to the hypotensive effects which have previously been reported^35^. Standardising this variable enables appropriate prioritisation of tasks during R-IRI surgery (placement of clamps at the correct time should be prioritised over closure) to achieve a more reproducible model. Standardisation causes a reduction in both GFR and variability, giving the potential to reduce the number of animals required to show therapeutic efficacy. It is well recognised that many factors including mouse strain, gender, age and temperature during surgery affect the degree of injury^11^. Given the number of variables that affect the injury level, refinement of the model by recognising and standardising these factors enables fewer animals to power a study.

### MSOT can be used to determine the function of an injured kidney after R-IRI

Previous publications have described measuring the delay between t_max_ in the renal cortex and pelvis^16^ using MSOT, and the area under the curve (AUC) after drug induced global renal injury^36^. Neither of these could be applied to the R-IRI model due to the difficulty of imaging both renal pelvises in one plane. This problem is compounded following surgery due to post- operative oedema and over-lying sutures. However, it was possible to accurately identify and image both renal cortices simultaneously after R-IRI, and obtain consistent t_max_ measurements.

Whilst it is not appropriate to use IRDye clearance to calculate GFR, due to its high levels of protein binding^16,25^, it can be used as a surrogate measure of renal function due to its sole excretion through the glomeruli. After assessment of IRDye PK, we found that t_max_ is the parameter that best distinguishes the injured from the uninjured kidney. Additionally, we determined that the injured kidney, similarly to the trajectory in the acute model, recovered by day 10 followed by prolongation of t_max_, indicating the progression from AKI to CKD. Whilst the development of CKD after AKI has been well described^37^, this is the first time that MSOT has been used to track CKD development longitudinally.

### Contralateral nephrectomy on day 14 after prolonged unilateral R-IRI produced a survivable, reproducible CKD model

Following the pilot experiment displaying proof of principle, refinement of the CKD model was undertaken by performing contralateral nephrectomy on day 14. While reduction in GFR persisted after contralateral nephrectomy, a slight non-significant upward trend was seen which probably represented the adaptation that has been described due to glomerular hyperfiltration^38^. However, the GFR reduction in week 6 remained significant compared to baseline levels. Development of this model demonstrates the utility of long-term accurate GFR monitoring, particularly when assessing the impact of therapeutic agents in an animal with predictable CKD.

In conclusion, the application of in-vivo non-invasive measures of renal function have enabled the refinement of models of both AKI and CKD after R-IRI. Transdermal GFR measurements correlate well with histological markers of kidney disease and enable the refinement and reduction of animals required for future studies of therapies in these models. MSOT is feasible in animals following R-IRI surgery and modelling of the clearance of IRDye using a tri-exponential curve and t_max_ enables reproducible monitoring of the function of the injured and uninjured kidney.

Application of the RIFLE criteria for GFR to pre-clinical studies enables the translation of results to the clinic, ensuring that therapeutic studies show clinically important differences.

## Author Contributions Statement

Funding for this work was secured by PM, BW, SK, PC, AO and RH. RH, PM, BW, SK, PC and AO designed the experiments. RH, LS and JS performed the animal studies including surgical intervention, measurements of FITC-Sinistrin and IRDye 800 clearance and analysis of weight change, serum creatinine, urea and Cystatin C. LR and RH performed the histological analysis. JB and GC performed the MSOT data analysis, RH, JB and GC undertook the statistical analysis. RH, JB, PM, BW and PC wrote the manuscript and generated the figures and all authors reviewed the manuscript prior to publication.

## Acknowledgements

We acknowledge funding from the Alder Hey Children’s Kidney Fund (RH, BW, PM, SK), Kidney Research UK (Fellowship awarded to RH, and Innovation grant awarded to PM, BW, LR, PK, AO) and UKRMP Safety and Efficacy Hub (PM, LS, JS, BW, LR). We thank the ‘Centre for Pre-Clinical Imaging’ at the University of Liverpool for facilitating these experiments and Dr M Garcia-Fiñana for her advice on experimental design and statistical analysis.

## Supplementary Information Contents

1. **Methods**
  a. **Anaesthetic and monitoring of animals**
  b. **Acute Kidney Injury (AKI) Model**
  c. **Refined AKI Model**
  d. **Multispectral Optoacoustic Tomography (MSOT) assessment of renal function**
  e. **Table 1: Key features of experimental series**
2. **Table 2: Serum Results – Initial and refined CKD model**
3. **Figures**
  1. **Modified operative steps of R-IRI**
  2. **Schematic of CKD Models of Injury**
  3. **Normality plot of baseline GFR and relationship to RIFLE criteria**
  4. **Weight change and mortality after AKI secondary to R-IRI**
  5. **Modelling of the clearance of IRDye800 using MSOT**
  6. **Cortical Collagen Deposition after CKD**

