## Supplementary Information for "Murine models of renal ischaemia reperfusion injury: An opportunity for refinement using non-invasive monitoring methods"

**Supplementary Information and Figures**

**Contents**

1. **Methods**
   a. **Anaesthetic and monitoring of animals**
   b. **Acute Kidney Injury Model**
   c. **Refined AKI Model**
   d. **Multispectral Optoacoustic Tomography (MSOT) assessment of renal function**
   e. **Table 1: Key features of experimental series**
2. **Table 2: Serum Results – Initial and refined CKD model**
3. **Figures**

**1. Modified operative steps of R-IRI**

**2. Schematic of CKD Models of Injury**

**3. Normality plot of baseline GFR and relationship to RIFLE criteria**

**4. Weight change and mortality after AKI secondary to R-IRI**

**5. Modelling of the clearance of IRDye800CW using MSOT**

**6. Cortical Collagen Deposition after CKD**

**1.a. Anaesthetic and monitoring of animals**

At the start of the surgical procedure animals weighed between 20.4 and 28.3g.

All surgical procedures were performed in the morning in an animal theatre. Oxygenated isoflurane was selected for anaesthesia due to its rapid onset and rapid recovery, along with the ability to be titrated to the animal’s response.

Animals were monitored throughout the procedure by the researcher and an animal technician. All animals were reviewed twice daily and more frequently if they showed any signs of distress.

**1.b. Acute Kidney Injury Model**

Bilateral renal pedicle clamping was performed at the age of 8-10 weeks. Anaesthesia was induced with inhaled oxygenated isoflurane (3%) in a box. The animal was transferred to a nose cone, the anaesthetic reduced to 1.75 ± 0.25% and received buprenorphine, baytrill and 0.5ml saline subcutaneously. A complete dorsal shave was performed and iodine was applied. The animal was transferred to a heat pad (Far Infrared Warming Pads, Physiosuite, Kent Scientific) with a rectal temperature probe to provide negative feedback. The body temperature was set to 37^0^C and the surgery was commenced after the temperature reached 36.5^0^C. A dorsal approach was taken^1^ to each kidney. The skin and then underlying muscle were incised and stretched open in the direction of fibres. The kidney was delivered using gentle pressure under the anterior abdomen and was then held using wide curved forceps. A pair of blunt dissecting forceps was used in the opposite hand to sweep the fascia and fatty tissue which overlies the hilum medially. The blunt forceps were then used to make a space above and below the renal vessels, removing as much of the perihilar fat as possible without causing bleeding. An atraumatic vascular clamp (Schwartz, Interfocus, Linton) was placed on the vessels using clip applying forceps. A timer was commenced for a pre-determined duration and the kidney was placed under the skin after confirming that the kidney was becoming uniformly dark. The same procedure was immediately performed on the contralateral kidney with the same duration of clamping. After the pre-determined duration of clamping, the clamp was removed, the kidney checked for re-perfusion and then replaced into the abdomen using a damp cotton-bud to gently push it back in. The muscle was closed using continuous 6/0 absorbable braided suture (CliniSorb, CliniSut sutures) and the skin with a horizontal mattress of the same suture. The animal was placed in a heat box to recover for 30 minutes at 32^0^C before being returned to their cage.

Surgery was performed on up to 3 mice at once with each mouse having its own heat pad and temperature probe. Mice were anaesthetised individually and then placed on a nose cone attached to an anaesthetic ‘splitter’.

3 groups of 6 mice underwent bilateral clamping for a duration of 25, 27.5 or 30 minutes. Data were collected on the duration of anaesthetic before the first clamp was applied and the duration between the second clamp’s removal and entering the heat box.

Whilst undertaking the initial bilateral R-IRI experiments, additional aspects of the experiment were observed and recorded whenever possible. These included (1) total anaesthetic time: from entry into the anaesthetic box to entry to heat box, (2) pre-clamping anaesthetic time: from entry into the anaesthetic box to application of the first clamp, (3) post-clamping anaesthetic time: from removal of the second clamp to entry into heat box, (4) experimental order, (5) heat pad.

**1.c. Refined AKI Model**

In addition to standardising the pre-anaesthetic duration, surgical loupes were worn to perform the procedure. An opened out vascular clamp was placed around the hilar vessels after their dissection to maintain the surgical plane created whilst the clamp was being prepared for application.

**1.d. Multispectral optoacoustic tomography (MSOT) assessment of renal function**

Multi-spectral optoacoustic tomography (MSOT) is an imaging system which causes excitation of endogenous and exogenous substances in the near-infrared range and through the process of thermo-elastic expansion gives a detectable acoustic wave^2^. IRDye800-CW is an organic flurophore which is renally excreted. It has maximum excitation at 775nm and has previously been shown to correlate well with renal function^3^.

Animals were anaesthetised with 3% oxygenated isoflurane which was reduced to 2% after anaesthesia was achieved. The animal was shaved on its dorsal and ventral sides of the abdomen and was depilated in the same area. A tail vein cannula was sited using a 30G broken needle inserted into 28G fine bore polyethylene tubing (Smiths, UK) which had been cut to a length of 32cm. A U-100 insulin syringe (BD Micro-Fine, Dickinson and Company, USA) was filled with 150 μl 0.9% saline, the pre-fixed needle was passed into the opposite end of the tubing and 50 μl saline was flushed through the tubing to prime it. The cannula was flushed with 100 μl 0.9% saline after placement and prior to imaging to confirm its position within the vein.

A flexible polyethylene membrane covered the MSOT frame and was covered with a thin layer of ultrasound gel in the area of the animal’s back. The animal was placed supine and the feet were secured. The animal was transferred into the MSOT with ongoing inhaled 2% oxygenated isoflurane and equilibrated to the water bath temperature of 34^0^C for 10 minutes. IRDye800CW (LI-COR, USA) was administered at a concentration of 0.1mg/ml, 220μl was prepared in a U-100 insulin syringe and was kept in the dark until required.

Imaging of the animal was taken in a single plane which best displayed both kidneys and renal pelvises. Images were acquired at 850nm and 775nm at a rate of 10 frames per second and averaging 20 consecutive frames for 3 minutes at which point the IRDye800CW was administered and the imaging continued for a further 20 minutes.

Images were reconstructed and underwent multispectral processing using the MSOT (iThera) software. The 850nm wavelength images were removed from the 775nm wavelength images after resolving all images for background signal. A region of interest was drawn around the area of highest intensity signal seen in the cortex of each kidney to determine the change in mean pixel intensity over time. The renal pelvis was not included in the analysis as the differing level of the two kidneys meant that both renal pelvises were rarely in view at the same time.

Previously, renal clearance has been characterised by two- or three-compartment models^4,5^. Here, we used MSOT intensity as a measure of IRDye800-CW concentration.

Pharmacokinetic (*PK*) profiles were assessed using the four summary measures: maximum peak of the curve (*c_max_),* the time at which *c_max_* occurs (*t_max_*), the terminal half-life (*t_50_*) and the area under the curve (*AUC*)^6^. A linear mixed effects model was fitted to the summary measures using R package NLME^7^. ‘Day’ and ‘injured versus healthy’ kidney were used as fixed effects to characterise the recovery of the kidney function over time and the subject was used as a random effect. The mixed effects models were used to assess if any of the summary measures could differentiate between injured and non-injured kidneys and were fitted using restricted maximum likelihood^8^.

**1.e. Key features of experimental series**

| **Experiment Description** | **Descriptor of groups** | **Animals per group**  **(n)** | **Analysed animals**  **(n)** | **Outcome Measures** | **Figures** |
| --- | --- | --- | --- | --- | --- |
| **AKI Model** | **Sham**  **25 minute**  **27.5 minute**  **30 minute** | **3**  **6**  **6**  **6** | **3**  **5**  **5**  **4** | **Mortality**  **Weight change**  **GFR**  **Serum analysis**  **PSR score** | **Supp F4B**  **Supp F4A**  **Figure 1A**  **Figure 2A,B,C**  **Figure 3A** |
| **Refined AKI Model** | **27.5 minute** | **6** | **6** | **GFR** | **Table 1** |
| **CKD Model** | **40 minute-Left** | **3** | **3** | **t_max_**  **GFR**  **Kidney weight**  **PSR score**  **Serum analysis** | **Figure 5**  **Figure 6A**  **Figure 6B**  **Figure 6C**  **Supp T2** |
| **Refined CKD Model** | **40 minute-Right** | **6** | **5** | **t_max_**  **GFR**  **Weight change**  **PSR score**  **Serum analysis** | **Figure 7A**  **Figure 7B**  **Figure 7C**  **Figure 7D**  **Supp F6**  **Supp T2** |

Supplementary Table 1: Key features of experimental series

2. **Serum Results**

|  | Initial CKD model | Refined CKD model |
| --- | --- | --- |
| Creatinine (mg/dL) | 1.87 (0.24) | 0.97 (0.23) |
| Urea (mg/dL) | 396.0 (228.2) | 179.3 (41.51) |
| Cystatin C (mg/L) | 1.27 (0.57) | 1.76 (0.48) |

Supplementary Table 2. Serum analysis in CKD Models. Initial CKD model n=3, Refined CKD model n=5. Results shown as Mean (Std Dev).

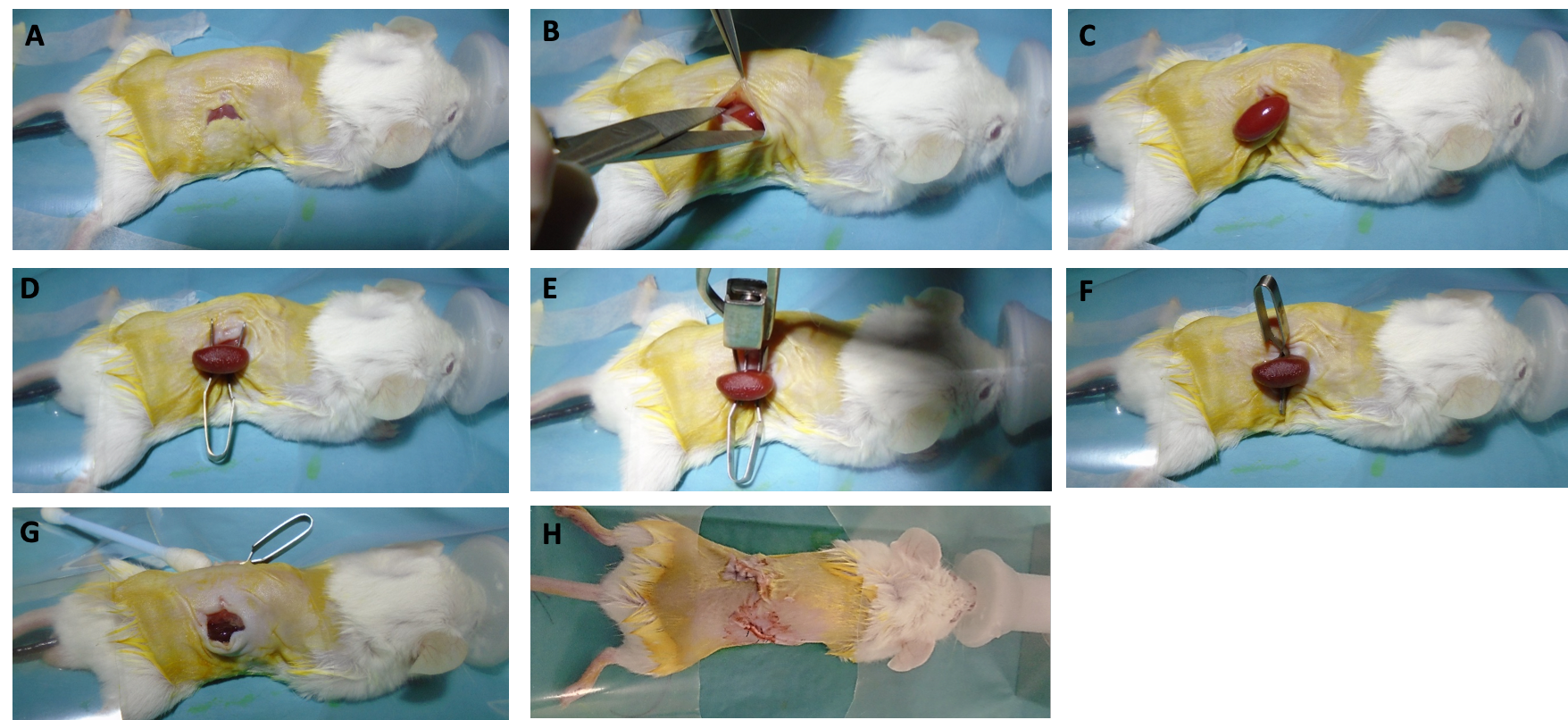

Supplementary Figure 1, Modified operative steps of R-IRI

A: Dorsal incision made over kidney

B: Muscle incised and split

C: Kidney delivered into wound

D: Hilar vessels cleared of fat by gently sweeping overlying fascia and fat supero-medially. Forceps and then opened vascular clamp placed around vessels

E: Vascular clamp placed around vessels

F: Uniform darkening of kidney seen. Kidney tucked under skin and operative steps repeated to contralateral side.

G: After clamping time complete, clamp removed and kidney observed for re-perfusion.

H: Continuous suture closure of muscle and interrupted horizontal mattress suture to skin.

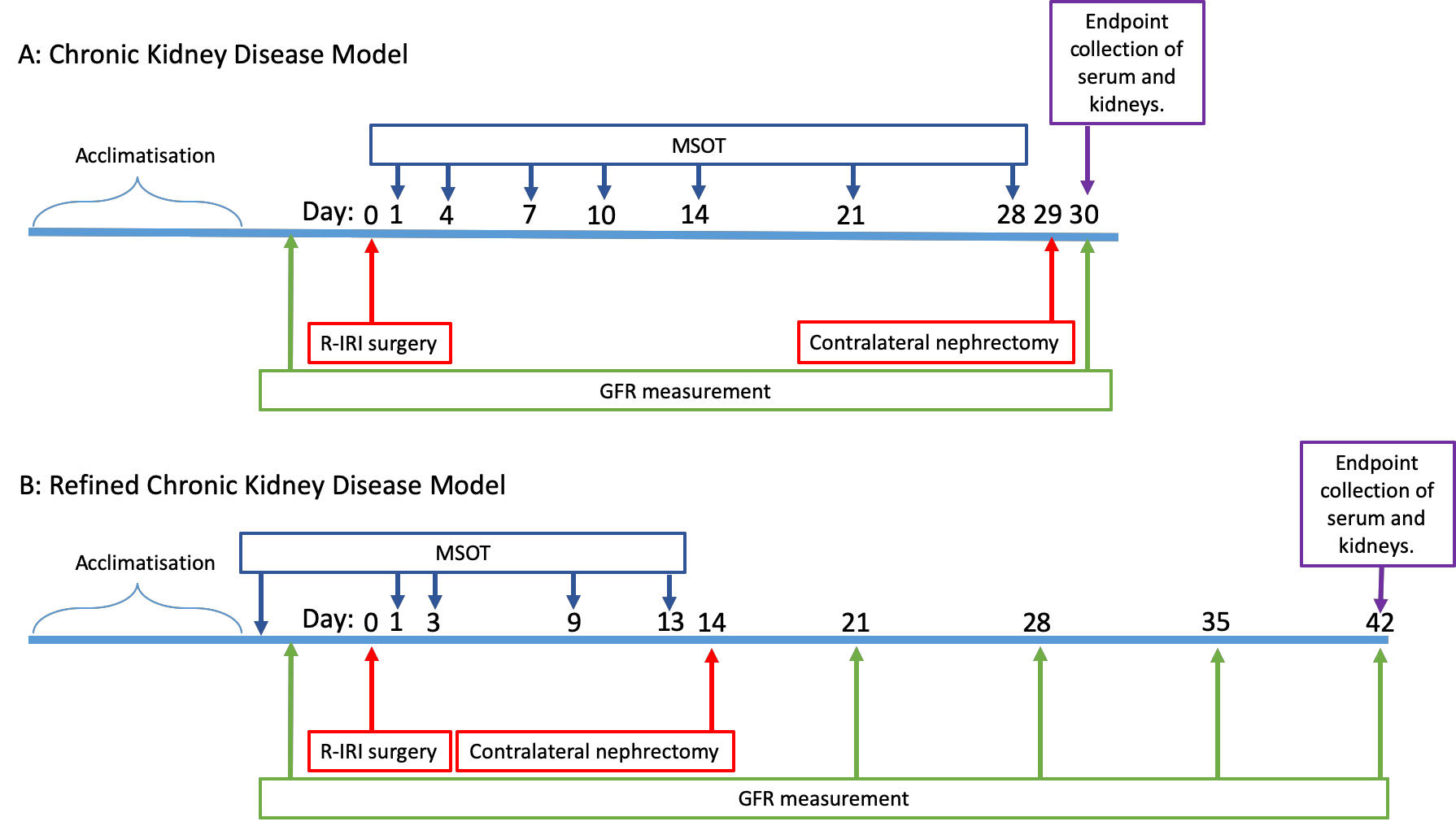

Supplementary Figure 2, Schematic of CKD models of injury.

A: Initial CKD model.

B: Refined CKD model

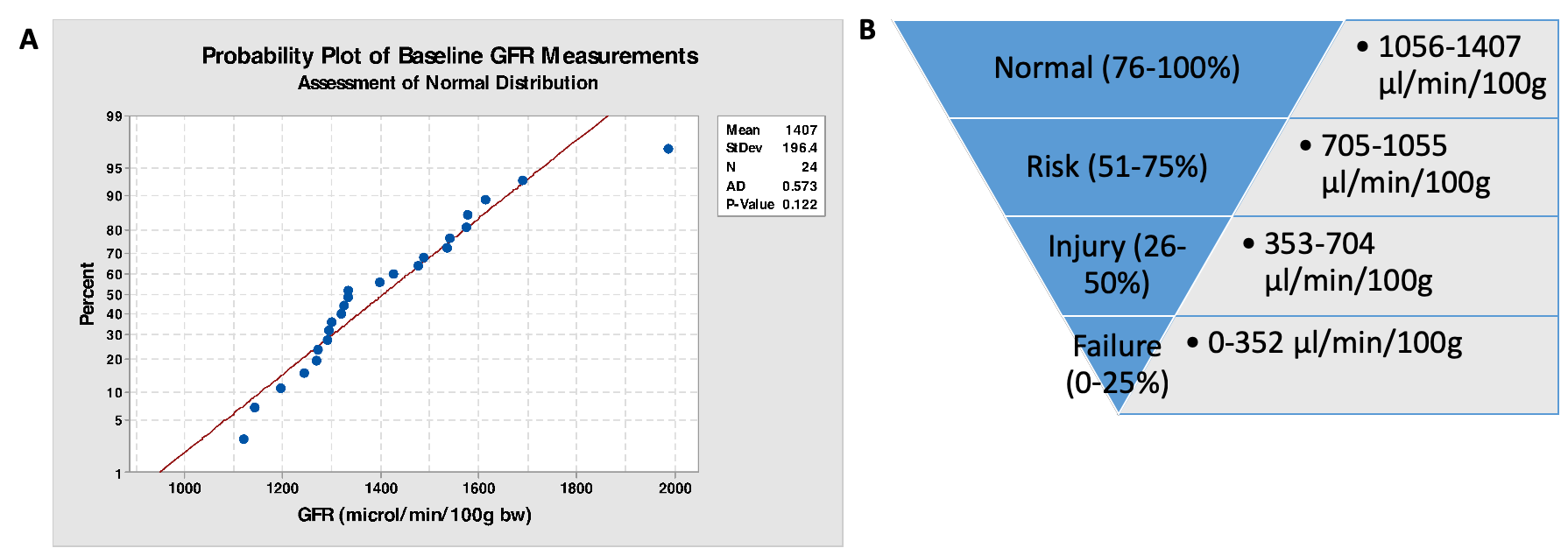

Supplementary Figure 3:

A: Normality plot of Glomerular Filtration Rate of uninjured male BALB/c mice at 7-9 weeks of age, showing that the probability that this is not a normal distribution is p 0.122. The red line depicts the expected spread of results if a normal distribution is present. The y-axis shows the percentage of values less that the GFR on the x axis.

B: Based on the Acute Dialysis Quality Initiative (ADQI) ‘RIFLE’ classification of GFR loss these values were applied to the data in this study.

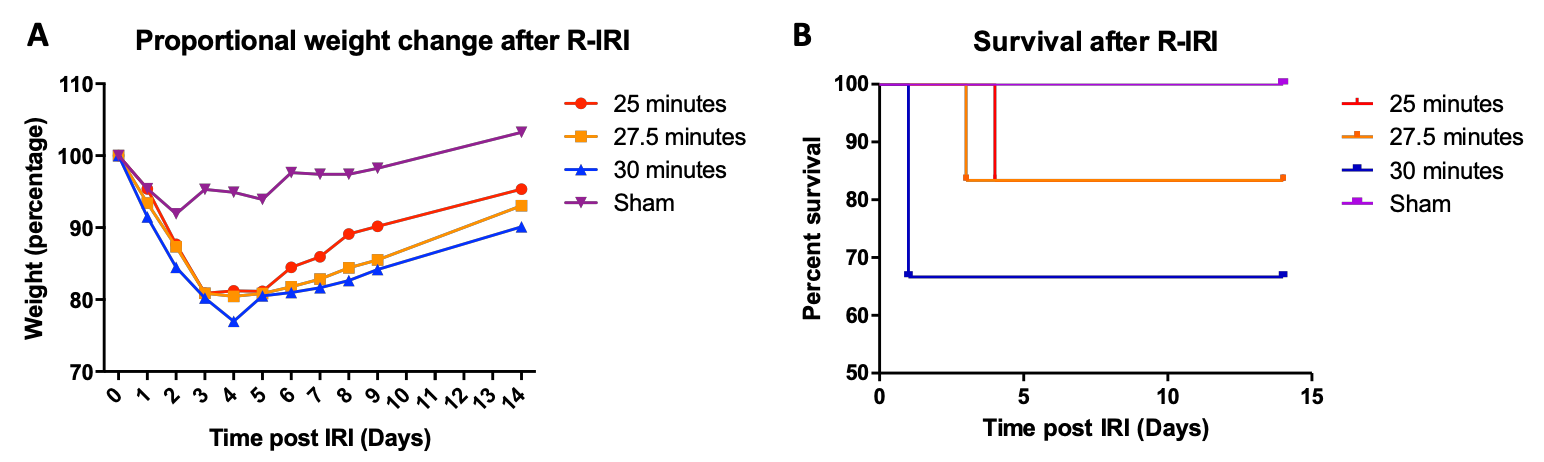

Supplementary Figure 4, A: Weight change after R-IRI in all animals who survived until the experimental endpoint of 14 days after R-IRI. Significant difference between sham and experimental groups on day 3 (25 minutes *, 27.5 minutes *, 30 minutes **), between sham and 30 minutes on day 4 and 6 *. B: Kaplan-Meier survival curve after renal ischaemia reperfusion injury (R-IRI). * p<0.05, ** p<0.01

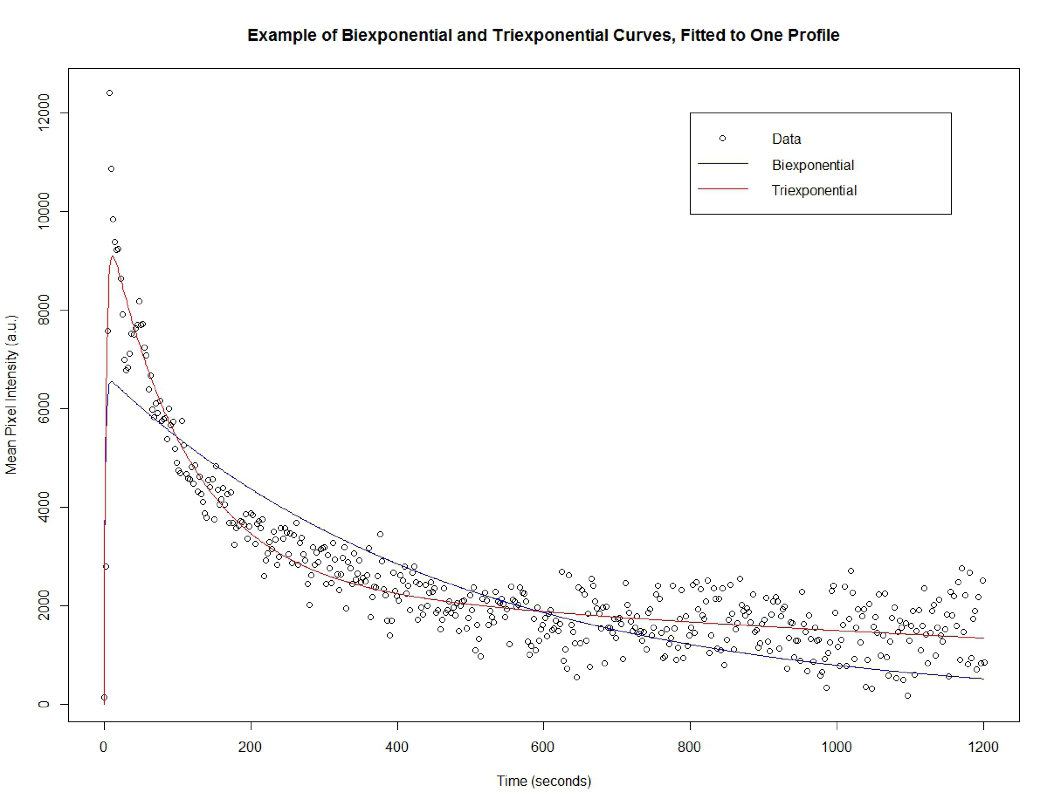

Supplementary Figure 5: modelling of the clearance of IRDye800 from the cortex of the kidney using biexponential and triexponential models.

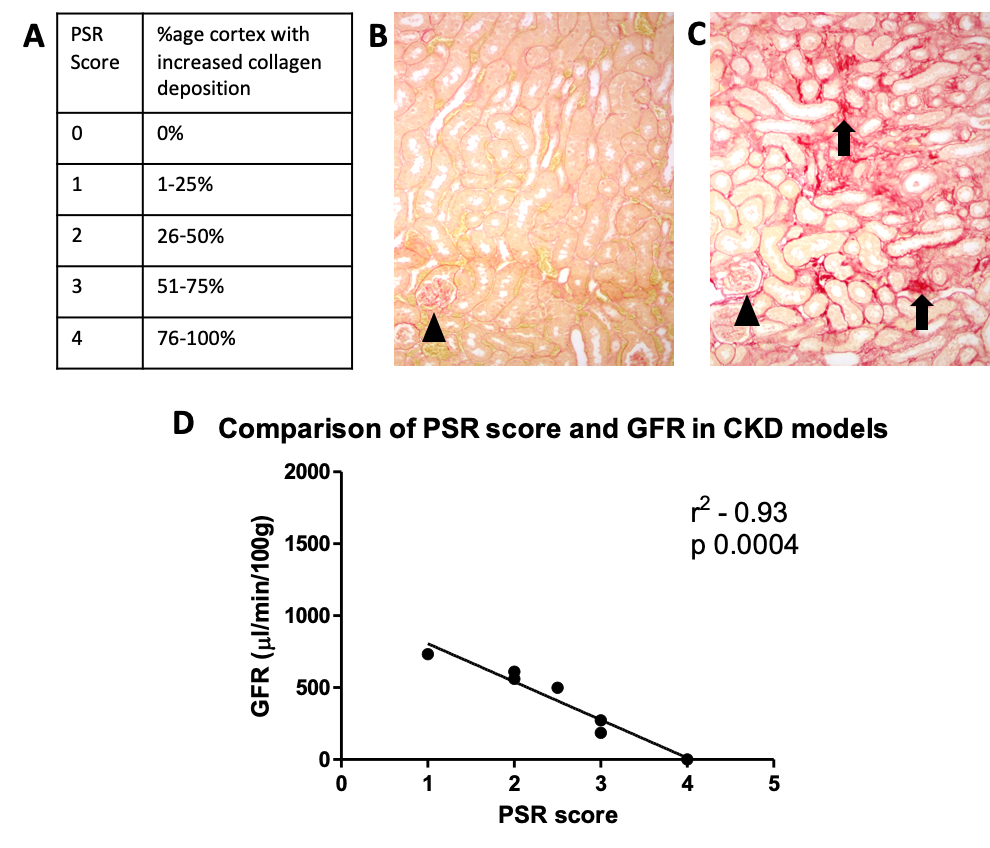

Supplementary Figure 6, Cortical collagen deposition

A: Scoring method for determining the percentage of the renal cortex which is affected by increased collagen deposition, as judged by Veterinary Histopathologist.

B & C: Picrosirius Red Staining (PSR) of renal cortex. Glomeruli marked with arrowhead, areas of increased collagen deposition marked with arrow. B shows no increase in collagen deposition, C shows areas of increased collagen deposition.

D: Correlation of PSR and endpoint GFR from the chronic kidney disease IRI models.

1	Skrypnyk, N., Harris, R. & Caestecker, M. d. Ischaemia-referfusion Model of Acute Kidney Injury and Post Injury Fibrosis in Mice. *JOVE* **e50495**, doi:10.3791/50495 (2013).

2	Taruttis, A., Morscher, S., Burton, N. C., Razansky, D. & Ntziachristos, V. Fast multispectral optoacoustic tomography (MSOT) for dynamic imaging of pharmacokinetics and biodistribution in multiple organs. *PLoS One* **7**, e30491, doi:10.1371/journal.pone.0030491 (2012).

3	Scarfe, L. *et al.* Measures of kidney function by minimally invasive techniques correlate with histological glomerular damage in SCID mice with adriamycin-induced nephropathy. *Sci Rep* **5**, 13601, doi:10.1038/srep13601 (2015).

4	Tofts, P. S., Cutajar, M., Mendichovszky, I. A., Peters, A. M. & Gordon, I. Precise measurement of renal filtration and vascular parameters using a two-compartment model for dynamic contrast-enhanced MRI of the kidney gives realistic normal values. *Eur Radiol* **22**, 1320-1330, doi:10.1007/s00330-012-2382-9 (2012).

5	Chen, B., Zhang, Y., Song, X., Wang, X. & Zhang, J. Quantitative Estimation of Renal Function with Dynamic Contrast-Enhanced MRI Using a Modified Two-Compartment Model. *PLOS ONE* **9**, e105087.

6	Urso, R., Blardi, P. & Giorgi, G. A short introduction to pharmacokinetics. *Eur Rev Med Pharmacol Sci* **6**, 33-44 (2002).

7	*The R Project for Statistical Computing*, <<www.R-project.org/>> (2018).

8	Morrell, H. Likelihood ratio testing of variance components in the linear mixed-effects model using restricted maximum likelihood. *Biometrics*, 1560-1568 (1998).
